# Design of a biosensor for direct visualisation of auxin

**DOI:** 10.1101/2020.01.19.911735

**Authors:** Ole Herud-Sikimic, Andre C. Stiel, Marina Ortega-Perez, Sooruban Shanmugaratnam, Birte Höcker, Gerd Jürgens

## Abstract

In plants, one of the most important regulative small molecules is indole-3-acetic acid (IAA) known as auxin. Its dynamic redistribution plays an essential role in virtually every aspect of plant life, ranging from cell shape and division to organogenesis and responses to light and gravity^1,2^. So far, the spatial and temporal distribution of auxin at cellular resolution could not be determined directly. Instead it has been inferred from visualisation of irreversible processes involving the endogenous auxin response machinery^3-7^. This detection system failed to record transient changes. Here we report on a genetically encoded biosensor for quantitative *in vivo* visualisation of auxin distributions. The sensor is based on the *E. coli* tryptophan repressor (TrpR)^8^ whose binding pocket was engineered for specific IAA binding and coupled to fluorescent proteins to employ FRET as readout. This sensor, unlike previous systems, enables direct monitoring of the fast uptake and clearance of auxin by individual cells in the plant as well as the graded spatial distribution along the root axis and its perturbation by transport inhibitors. Thus, our auxin sensor enables mapping of auxin concentrations at (sub)cellular resolution and their changes in time and space during plant life.

---

The tryptophan-derived metabolite indole-3-acetic acid (IAA; *vulgo* auxin) triggers a multitude of developmental processes and responses to environmental cues, thus conditioning plant life^1,2^. Much progress has been made in the past two decades towards mechanistic understanding of the nuclear events turning auxin perception into transcriptional responses^9-11^. Other studies have addressed the basic machinery of polar and non-vectorial release of auxin from the cell within a tissue context through the action of PINFORMED efflux transporters and ABCB transporters, resulting in various computer models of how canalised auxin flow mediates developmental or physiological processes^12-14^. In contrast, virtually nothing is known about the actual distribution of auxin in developing or growing tissues at single-cell resolution because of technical limitations (reviewed by ref. 15). Plant biologists could only use proxies such as auxin-dependent reporter gene expression to visualise auxin distribution (DR5::GUS^3^; DR5::ER-GFP^4^; DR5::NLS-3xGFP^5^); this indirect approach is characterised by latencies and possible non-linearities. More recently, IAA levels have been inferred from auxin-dependent degradation and thus signal reduction of fluorescent proteins linked to domain II of an inhibitory IAA protein^6,7^. A major caveat of these approaches is their irreversibility precluding visualisation of auxin transients.

In contrast, the ideal auxin sensor should have the following features: (1) Physical interaction with auxin should elicit a quantitative fluorescent signal in a reversible manner so that the rise and fall of auxin concentration can be monitored. (2) The sensitivity should be high enough so that a dynamic image of auxin distribution over time can be generated. (3) The sensor should be targeted to different subcellular compartments as well as to the extracellular space and thus to locations that are out of reach for the conventional proxies, which rely on gene expression or protein degradation. (4) Components of plant metabolism or regulation should not be used to reduce the likelihood of interfering with auxin response.

With these boundary conditions in mind, we have developed a genetically encoded, fully reversible biosensor for *in vivo* imaging of auxin gradients with high spatial and temporal resolution, starting from the tryptophan repressor (TrpR), a bacterial tryptophan-binding protein. Auxin bears a high resemblance to tryptophan (TRP); chemically, an amino acid group is exchanged by a carboxyl group that is connected via a carbon to the common indole ring (Fig. 1a). The dimeric TrpR undergoes a conformational change upon binding TRP that enhances its affinity for the operator DNA of the TRP biosynthesis operon^16,17^. Fluorescent proteins fused to TrpR relay the conformational change and translate it into a FRET signal, which is a convenient readout for *in vivo* measurements (Fig. 1b)^18^. Furthermore, TrpR exhibits already low affinity towards IAA^8^ (Fig. 1d). This makes TrpR a perfect starting point for developing an auxin-specific genetically encoded FRET biosensor^19^. Our design efforts were aimed at improving affinity and specificity of IAA binding while abolishing TRP binding. We assumed a comparable binding mode for the indole ring of both ligands and focused our design on TrpR residues in the vicinity of the TRP amino group (Fig. 1c), aiming to improve the specificity for the IAA carboxyl group. This selection was later expanded to adjacent residues. Altogether about 2,000 variants were generated in successive rounds of saturation mutagenesis and screened for increased FRET signal upon IAA addition (Fig. 1d and Extended data Fig. S1). Improved variants were regularly checked for specificity with a library of substances similar to IAA and reportedly present in *Arabidopsis* (Extended data Table S1). To confirm improvements in binding affinity, selected TrpR variants were purified and analysed by isothermal titration calorimetry (ITC, Extended data Table S2). Furthermore, the structures of several variants were elucidated by X-ray crystallography to guide site-directed mutagenesis (Extended data Table S3).

**Figure 1.**
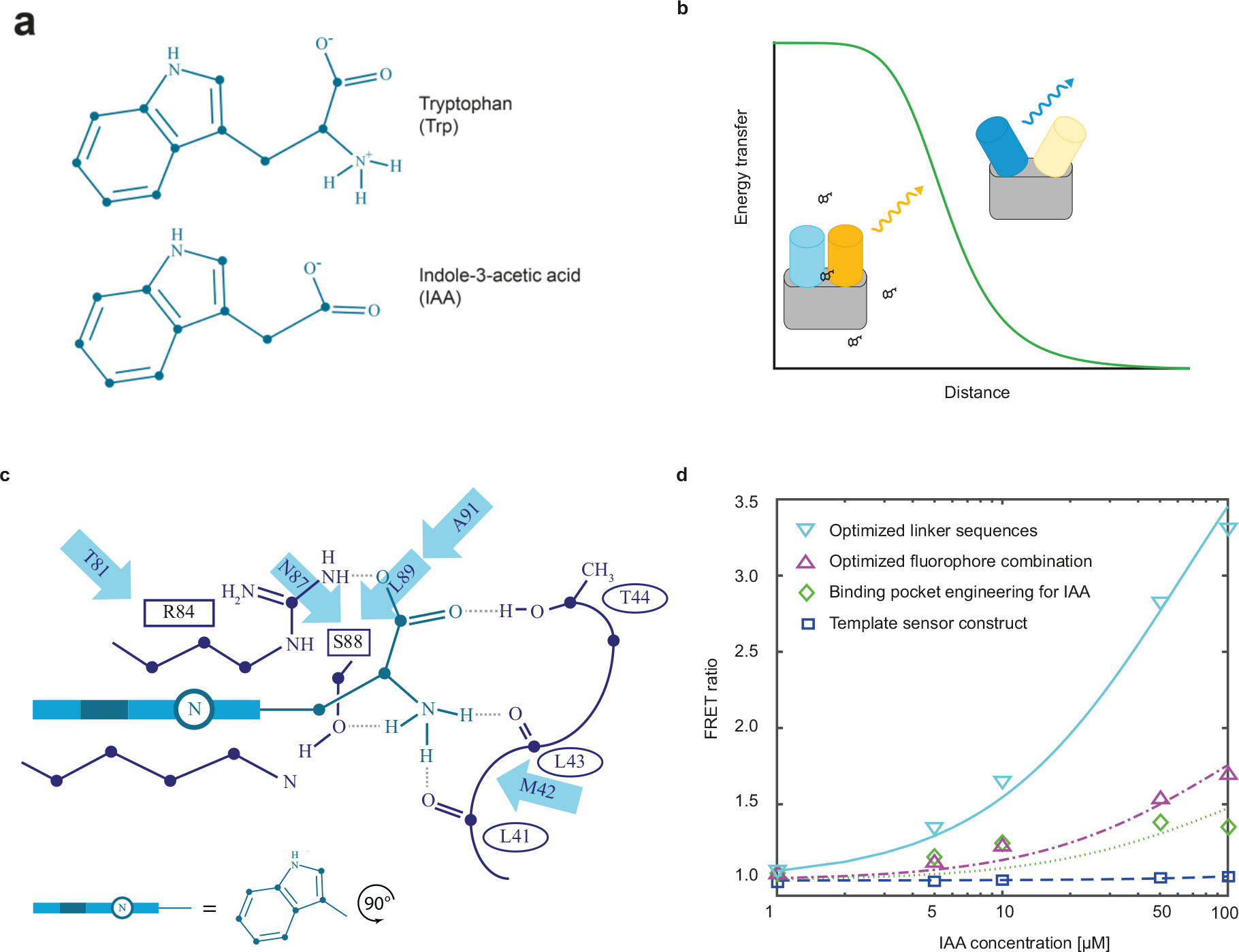
Summary of the design process. **a**, Chemical structures of TRP and IAA. **b**, Principle of the sensor design. In the presence of IAA, the fluorophores are close to each other and energy is transmitted (E_FRET_). In its absence, the distance increases and less energy is transmitted. **c**, Structure of the binding pocket of TrpR (modified from ref. 8). Interactions with the side chains of R84 and S88 as well as the backbone carbonyl groups of L41, L43, and T44 of the second chain are shown explicitly. Residues mutated in this study are indicated as arrows. **d**, FRET ratio change plotted against IAA concentration and contributions of the individual steps to the final sensor. Template sensor construct, TrpR wt – eCFP – Venus (blue); Engineered binding pocket for IAA, TrpR-M42F-T44L-T81M-N87G-S88Y – eCFP – Venus (green); Optimised fluorophore combination, TrpR-M42F-T44L-T81M-N87G-S88Y - mNeongreen – Aquamarine (purple); Linker optimised, TrpR-M42F-T44L-T81M-N87G-S88Y - mNeongreen - Aquamarine - Linker optimised (light blue).

Our structural analysis showed that in comparison to TRP binding (Fig. 2a), IAA is flipped by 180° with the carboxyl group facing the opening of the TrpR binding pocket (Fig. 2b). While TRP is anchored by interactions to the protein surrounding, IAA binding shows no such stabilisation, which was reflected in poor affinity (Extended data Table S2). In engineering the auxin sensor, we identified variants that stabilise and favour this IAA binding mode. Foremost, mutation S88Y entirely blocks TRP binding with its bulky sidechain while simultaneously favouring IAA through interaction of its carboxyl with the R84 guanidino and Y88 hydroxyl group, respectively (Fig. 2c). Further improvement of IAA affinity was achieved by optimising hydrophobic interactions of the indole ring with the binding pocket, i.e. with mutations T44L and T81M that both contribute to the final sensor (Fig. 2d). During the engineering process, we also monitored binding to possibly competing IAA-related plant compounds such as indole-3 acetonitrile (IAN) (Extended data Table S2). The modes of IAN and IAA binding are strikingly similar (Extended data Fig. S2a and S2b). Only few mutations like N87G exert discriminating effects mainly via small changes in the positioning of Y88 (Extended data Fig. S2c). Finally, we identified mutations that have no favourable effect on IAA affinity but improve the FRET readout most likely through small changes in packing and thus orientation of the attached fluorescent proteins (FPs) (Extended data Fig. S2d). In further steps, we optimised fluorophores as well as linker combinations (Extended data Fig. S3) to yield our final sensor AuxSen (mNeonGreen-TrpR-Aquamarine-TrpR, with TrpR being TrpR-M42F-T44L-T81M-N87G-S88Y, Fig. 1d and Fig. 2d).

**Figure 2.**
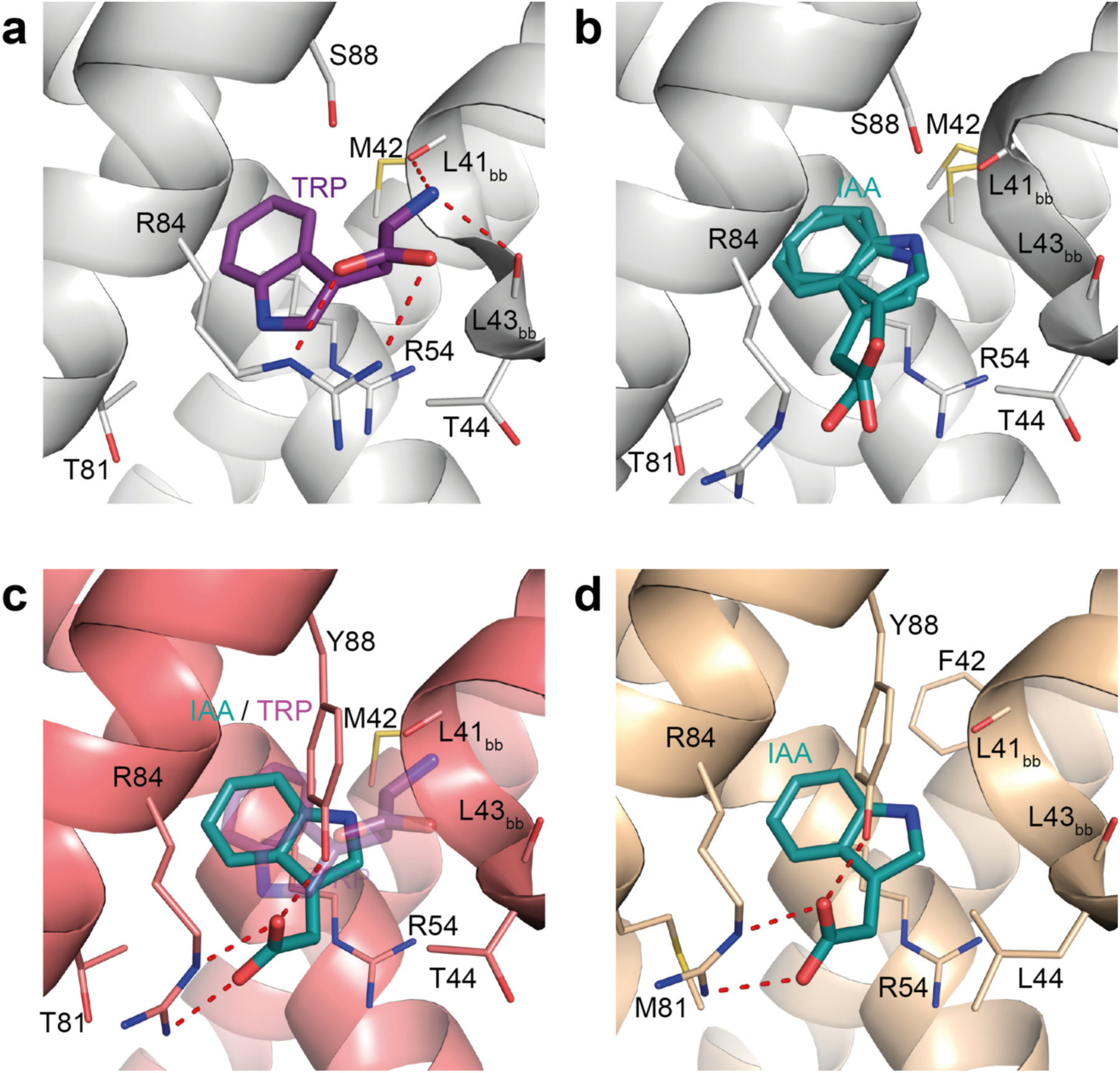
Structure of AuxSen and critical steps in the engineering process. **a**, Structure of TrpR-wt bound to the native ligand TRP (lilac, pdb-id 1ZT9) and **b**, bound to the design-target IAA (green). IAA shows a ligand position turned by 180° compared to TRP. Due to a lack of stabilisation in TrpR, IAA shows conformational freedom; two alternative conformations are shown. **c**, The mutation S88Y sterically precludes the position of TRP (transparent lilac) while favouring the position of IAA. **d**, Structure of the final AuxSen variant (TrpR-M42F-T44L-T81M-N87G-S88Y) bound to IAA. The ligand is firmly packed in the enhanced hydrophobic pocket and anchored to R84 as well as Y88, resulting in a high IAA affinity. All structures are superimposed on the Cα of residue 20-60 of both chains. Subscript “bb” labels residues whose backbone atoms interact with tryptophan in the native complex.

*In vitro*, the FRET ratio of AuxSen changed by a factor of three upon treatment with 50 µM IAA, which is in the range of cellular auxin concentrations^20^. The signal is stable at the cytosolic pH, resistant to reducing and oxidising environments and all tested salt ions (Extended data Fig. S4, Supplementary discussion). Specificity of AuxSen for IAA was tested against other indole derivatives reportedly present in *Arabidopsis* (Extended data Table S1). AuxSen has clearly the highest affinity for IAA, a few other compounds show a response but bind about one order of magnitude less well (Extended data Fig. S5). Of those compounds, only IAN is present in higher amount in plants (Supplementary discussion). However, roots show a growth response to treatment with IAN, and modelling suggests that the native IAA receptor SCF^TIR1^ could bind IAN^21^. Thus, it seems likely that IAN is sequestered and would not interfere with auxin sensing in the plant.

As a first step to confirm the functionality of the sensor *in vivo*, we expressed a nuclear-localised version of the sensor transiently under the control of the strong viral *35S* promoter in protoplasts^22^. The FRET ratio increased with the auxin level in the medium (Fig 3a and Extended data Fig. S6). The FRET ratio increasingly differed between cells at higher auxin concentrations, suggesting that IAA was not taken up by all cells with equal efficiency (Fig. 3a). We also examined whether the sensor can report auxin in the endoplasmic reticulum (ER) because compartmentalisation might be an ancient mechanism of auxin homeostasis^23^. The ER-targeted sensor responded positively when protoplasts were incubated in auxin-containing medium (Fig. 3b).

**Figure 3.**
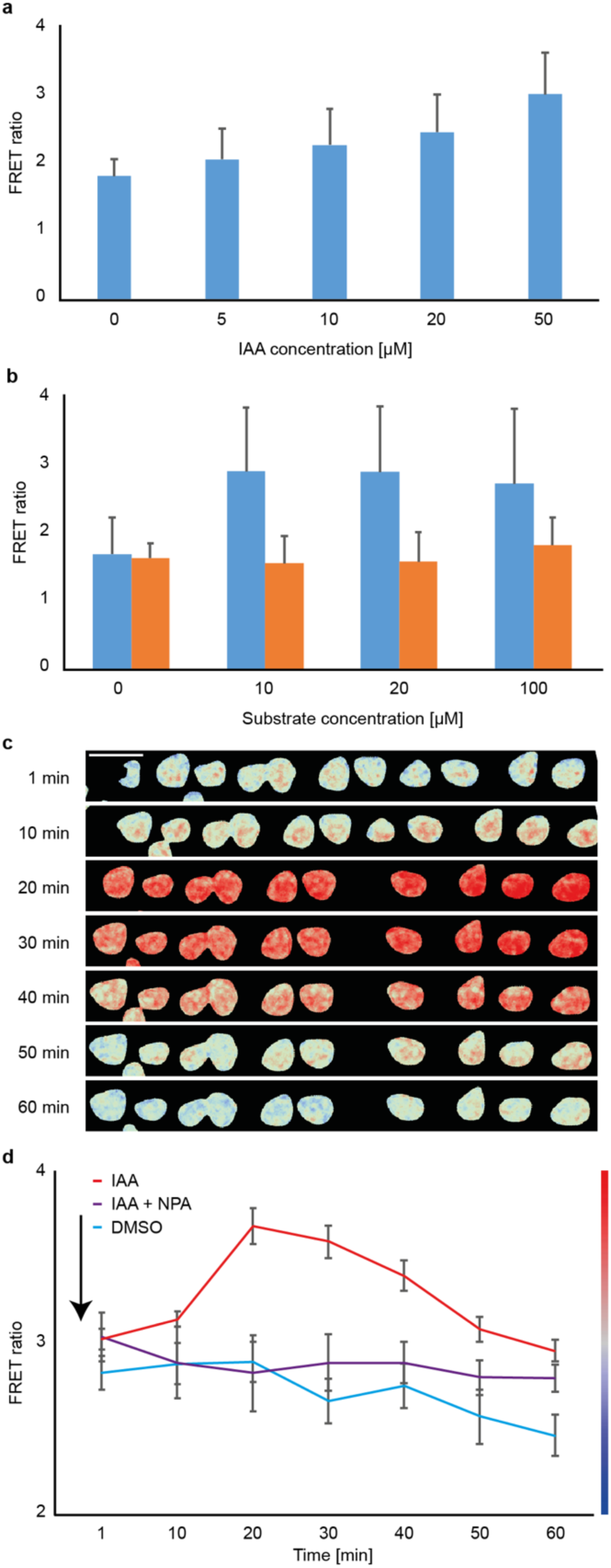
FRET ratio of auxin sensor in response to auxin treatment. **a–b**, Protoplasts were exposed to rising auxin concentrations in the medium: **a**, Nuclear-localized (NLS) auxin sensor; **b**, ER-targeted auxin sensor in the presence of IAA (blue) or the synthetic auxin 2,4-D (orange). **c–d**, Transient FRET ratio change of NLS auxin sensor in sub-epidermal cells of seedling roots treated with (**c**) 10 µM IAA; colour code indicated on the right in (**d**). **d**, Average FRET ratio of the 10 nuclei shown in (**c**) (IAA, red line) in comparison to control (DMSO, blue line) and a root pre-treated with NPA (IAA + NPA, purple line). The arrow indicates the start of IAA treatment. Error bars represent standard deviation.

To assess the functionality of AuxSen *in planta*, we initially generated lines expressing AuxSen under a variety of promoters. Overall, the *ELONGATION FACTOR 1a* promoter (*pEF-1a*)^24^ performed best. We checked its activity by driving the expression of β-glucuronidase (GUS), mNeonGreen and AuxSen. The promoter was particularly active in the root tip; weak expression was also detectable in cotyledons and shoot apical meristem (Extended data Fig. S7). AuxSen expression did not interfere with auxin signalling, as exemplified by root growth (Extended data Fig. S7e). Nevertheless, we observed silencing of the sensor in later generations, similar to other plant biosensors^25^. Heterozygous lines appeared stable and were used for subsequent experiments. To examine the response of AuxSen to auxin *in planta*, we treated seedlings with 10 µM IAA and recorded the FRET signal over time. After ten minutes, the AuxSen signal increased in cells of the sub-epidermal layer, reached a maximum after 20 minutes and then decreased slowly to background levels; in contrast, treatment with the solvent DMSO did not induce any response (Fig. 3c, d; Extended data Fig. S7f-l). This transient signal demonstrated the value of a sensor that binds auxin reversibly. Although traditional reporter systems can detect the response to auxin uptake^6^, the irreversibility of reporter translation or degradation obscures the transient nature of the auxin response.

We used a two-component expression system (*RPS5A::Gal4-GR X UAS::AuxSen*) to visualise the spatial distribution of endogenous auxin within the seedling root. The strong ubiquitously active promoter *RPS5A* drove expression of the yeast Gal4p transcription factor whose nuclear uptake was induced by dexamethasone (DEX), resulting in the expression of both the nuclear-localised auxin sensor and the nuclear-localised expression-control marker UAS::NLS:tdTomato. After DEX induction overnight, the FRET ratio steadily increased from the top end to the root tip (Fig. 4a). Interfering with auxin transport by incubating the seedlings in 50 µM NPA, which purportedly inhibits auxin transport^26,27^, altered the spatial distribution of the FRET signal. Specifically, the high end of auxin accumulation at the root tip was increased to give a peak at the expense of the adjacent region whereas the top end of the root was cleared of auxin (Fig. 4b, c). FRET ratios were significantly increased between 110 and 140 µm from the tip for NPA-treated roots compared to untreated controls (n_t_ = 11, n_c_ = 8; *t*-test, p<0.05). This change suggested not only auxin transport from the top end but also local auxin biosynthesis at the root tip, which normally appears to be masked by ongoing auxin transport both downward and upward^28^. We also treated seedlings with the fungal toxin brefeldin A (BFA), which interferes with auxin transport by inhibiting the BFA-sensitive ARF-GEF GNOM required for polar recycling of the auxin efflux transporter PIN1^29^. BFA treatment caused a comparable redistribution of auxin along the root length as observed before in NPA-treated roots (Fig. 4f, compare with Fig 4c). FRET ratios were significantly increased between 80 and 120 µm from the tip for BFA-treated roots compared to untreated controls (n_t_ = 13, n_c_ = 13; *t*-test, p<0.05). In conclusion, our auxin sensor responds to perturbation of endogenous auxin distribution caused by differently acting transport inhibitors in a consistent manner. This underlines the specificity of auxin detection of AuxSen.

**Figure 4.**
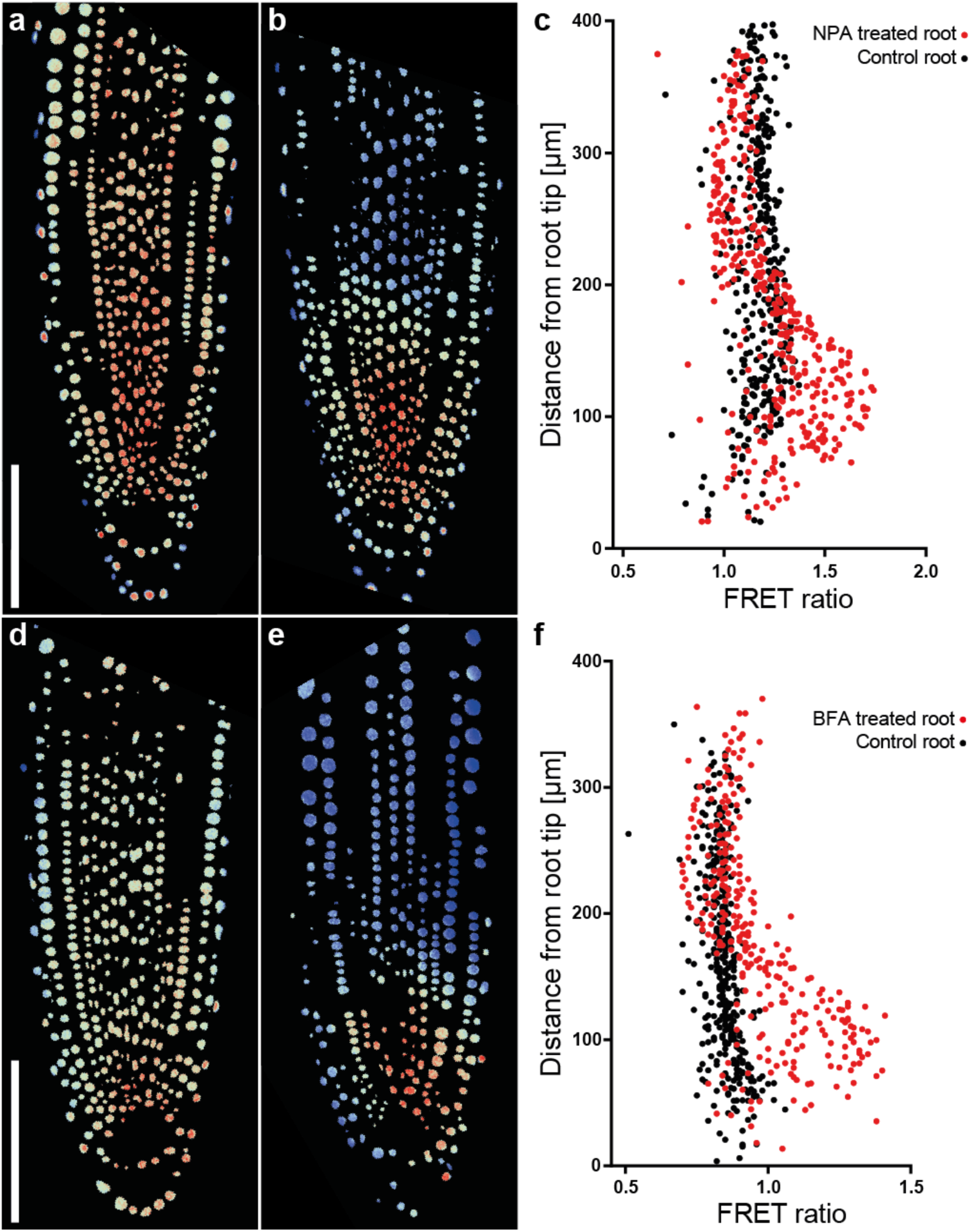
FRET ratio of auxin sensor in response to redistribution of endogenous auxin. **a–c**, Primary seedling root treated with (**a**) DMSO (mock treatment / control), or (**b**) auxin transport inhibitor NPA (50 µM for 24 hours); (**c**) quantitative FRET ratio distribution along the root (black, control; red, NPA-treated). Each dot represents a nucleus. **d-f**, Primary seedling root treated with (**d**) DMSO (mock treatment / control), or (**e**) membrane-trafficking inhibitor brefeldin A (BFA; 25 µM for 24 hours); (**f**) quantitative FRET ratio distribution along the root (black, control; red, BFA-treated). Each dot represents a nucleus.

Our semi-rational design approach has yielded a novel sensor for the pervasive signalling molecule auxin in plant development and physiology. Starting from a tryptophan sensor, we optimised affinity and specificity of small-molecule binding and improved signal intensity through the choice of FRET pair and linker optimisation. Our results provide a proof of principle that the new detection system can visualise the dynamic redistribution of auxin as well as subcellular and extracellular pools of auxin, which cannot be done with the reporters currently used. Furthermore, AuxSen enables changes in auxin distribution to be distinguished from changes in auxin response, which is a prerequisite for dissecting the complex regulatory network underlying the biological effects of this major signalling molecule in plant growth and development.

## METHODS SUMMARY

All variants were generated by site-directed mutagenesis and expressed in *E. coli* BLR(DE3). For initial screening, proteins were expressed in *E. coli* grown for 3 days at room temperature on plates in the dark, collected in MOPS buffer, sonicated and centrifuged. The supernatant was used to measure the FRET ratio upon addition of increasing amounts of the substrate with a Tecan plate reader. Confirmation screening was done with protein extract collected with His spin tubes (GE) according to the manufactures description. The buffer was exchanged against 20 mM MOPS with Illustra NAP-25 Columns.

Proteins for ITC and structure determination were expressed in *E. coli* BL21(DE3) in shaking flasks and purified using standard affinity and size exclusion chromatography. Crystals were obtained by vapour diffusion and data were collected at synchrotron beamlines by single-crystal X-ray diffraction. Structures were solved by molecular replacement and deposited with the PDB under the following accession codes: 6EJW, 6EJZ, 6ENI, 6EKP, 6ENN, 6ELB, 6ELF, 6ELG. ITC was performed using a VP-ITC (MicroCal) instrument with the ligand titrated to a protein solution of fixed concentration.

Protoplasts were transfected with 10 µg of the sensor construct and imaged with LSM780 (Zeiss) in K3 medium. For live imaging, seedling roots were incubated in PBS +15% glycerol, for steady-state images roots were incubated in PP11.

## Acknowledgements

We thank Addgene for distributing plasmids donated by Wolf Frommer, Fabienne Merola, Kurt Beam, Kurt Thorn and Oliver Griesbeck, Megan Sawchuk for the *pDR5rev::mRFP1er* seeds, Helen Schäfer, Anja Holz and Ana-Cristina Barragan-Lopez for technical assistance, and Mireille Belkacemi for sequencing of the variants. We further thank the beamline staff at the Swiss Light Source and at BESSY for support. B.H. acknowledges the financial support and the allocation of synchrotron beam time by HZB as well as funding by the Deutsche Forschungsgemeinschaft (grant HO 4022/2-3).

## Author Contributions

OHS, AS, BH and GJ conceived the idea and designed the experiments, OHS, AS, SS and MOP performed the experiments, OHS, AS, MOP, BH and GJ wrote the manuscript with input from all authors.

## Author Information

Correspondence and requests for materials should be addressed to Gerd Jürgens or Birte Höcker.

