## Supplementary material for "Design of a biosensor for direct visualisation of auxin": Suppl. Material

**Full Methods**

### *Cloning of in vitro auxin sensors*

The TrpR sensors Trp-CTY and Trp-CTYT were gifts from Wolf Frommer (Addgene plasmids #13533 and #13534). The fluorophores tested were also kindly donated: Aquamarine by Fabienne Merola (Addgene plasmid #42888), Clover and mRuby2 by Kurt Beam (Addgene plasmid #49089), mKO1 by Kurt Thorn (Addgene plasmid #44642), mCherry by Martin Bayer[30]. eGFP was amplified from pGIIK NLS::3xEGFP[31]. mNeonGreen, mWasabi and mTFP1 were purchased from Allele Biotechnology & Pharmaceuticals, Inc. San Diego, CA, USA, mKate2, and TagRFP from Evrogen, Moscow, Russia.

For the initial screening, we first used the Trp-CTY sensor and mutated the residues T44, S88 individually to all possible amino acids. To this end, we generated primers with 15-16 bp overlaps around the exchanged amino acid codons. We first mutated the amino acids sequence randomly with a degenerate primer and screened 96 clones. Variants not found were then generated by targeted mutagenesis. E.g. to generate all T44 variants we first used the primers CCTGATGCTGNNNCCAGATGAGCGCG and CGCGCTCATCTGGNNNCAGCATCAGG, to generate the missing T44C variant we then used the primers CCTGATGCTGtgtCCAGATGAGCGCG and CGCGCTCATCTGGacaCAGCATCAGG. The most promising candidates were introduced by targeted mutagenesis into Trp-CTYT and TrpR without FPs, which were analyzed by ITC and structural studies. Trp-CTYT with mutations T44L and S88Y was generated in 2 steps: we first used the primers CCTGATGCTGctaCCAGATGAGCGCG and CGCGCTCATCTGGtagCAGCATCAGG to generate Trp-CTYT with T44L and afterwards GATTACGCGTGGATCTAACtac-CTGAAAGCCGCGCCC and GGGCGCGGCTTTCAGgtaGTTAGATCCACGCGTAATC to generate Trp-CTYL with T44L and S88Y. To produce the recombinant proteins we cloned the TrpR domain into the pET21 expression vector and generated the variants by targeted mutagenesis. This procedure was repeated with the most promising variants after each round. The primer sequences are available upon request.

The final variant was then codon-optimised for *Arabidopsis* and synthesised by Thermo Fisher Scientific GENEART GmbH, Regensburg, Germany. To allow an easy exchange of the fluorophores we added restriction enzyme sites at the ends: *Bam*H1 and *Xho*I around the first fluorophore and *Apa*I and *Hind*III around the second. All fluorescent proteins were tested in the fluorophore I – TrpR – fluorophore II – TrpR configuration. Having identified Aquamarine and mNeongreen as the optimal pair, we introduced all final binding pocket variants into the backbone by site-directed mutagenesis.

The *pDR5rev::mRFP1er* line was a gift from Megan Sawchuk[32].

#### *Ligands used for screening and testing*

IAA, TRP, IAN, indole-3-carboxaldehyde, indole, indole-3-acetyl alanine, indole-3-acetyl aspartic acid, indole-3-acetamide, indole-3-ethanol, L- kynurenine, 2-oxindole-3-acetic acid, phenylalanine, picloram, tryptamine, $(NH_4)_2SO_4$, $CaCl_2$, DTT, $H_2O_2$, $NH_4NO_3$, $KNO_3$, and yucasin were purchased from SIGMA-ALDRICH, Saint Louis, USA; 4-hydroxyindole-3-carbaldehyde and 4-hydroxyindole-3-carbaldehyde from Santa Cruz Biotechnology, Inc., Santa Cruz, USA; 1-naphthaleneacetic acid, KCl, and DMSO from Carl-Roth GmbH, Karlsruhe, Germany; 2,4-dichlorophenoxyacetic acid (2,4-D) from Alfa Aesar, Karlsruhe, Germany; indole-3-acetyl glucose from TRC, North York, Canada; indole-3-butyric acid from SERVA, Heidelberg, Germany; NaCl from Merk, Darmstadt, Germany; NPA from SUPELCO, Bellefonte, USA; brefeldin A (BFA) from Thermo Fisher Scientific.

#### *Mutagenesis*

Trp-CTY variants were generated by site-directed mutagenesis with degenerate or specific oligonucleotides purchased from SIGMA-ALDRICH, Saint Louis, USA. Amplification was done with *Pfu* Polymerase (Thermo Fisher Scientific).

More than thousand oligonucleotides were used; the sequences and resulting vector maps are available upon request. Each variant was sequenced and screened in crude extract of sonicated bacteria for IAA binding, promising candidates were confirmed as purified proteins. The linkers were generated by site-directed mutagenesis, including linkers with 15-16 bp overlap and 3 to 9 degenerated nucleotides in the middle. To generate linkers shorter than the original ones, parts were deleted whereas for longer linkers, a fixed sequence was inserted into the middle to reduce the risk of generating stop codons by having too many degenerate nucleotides in the sequence.

#### *Protein expression and purification for screening*

For protein expression, bacteria were grown in the dark on plates with LB-Agar supplemented with Ampicillin for 3 days at room temperature. To measure crude extracts, we resuspended the bacteria in 20 mM MOPS pH 7.2, sonicated the suspension with a MS 73 probe (Bandelin, Berlin, Germany) and centrifuged the sample with a tabletop centrifuge (Eppendorf, Hamburg, Germany). Protein extraction was performed with His Spin trap columns according to the manufacturer´s instructions (GE Healthcare, Buckinghamshire, UK). We resuspended the bacteria of one petri dish in 2 ml binding buffer and sonicated with a MS 73 probe. Buffer exchange was performed with illustra NAP-25 Columns (GE Healthcare, Buckinghamshire, UK). All measurements were performed in 20 mM MOPS (SIGMA-ALDRICH, Saint Louis, USA) with an Infinite F200 plate reader (Tecan, Männedorf, Switzerland).

#### *Ligands*

Indole-3-acetic acid, indole-3-acetonitrile, and L-tryptophan used for crystallization and ITC were obtained from Sigma-Aldrich. All ligands were dissolved in 50mM Tris / 300mM NaCl pH8 buffer containing 1% dimethyl-sulfoxide (DMSO) if necessary.

#### *Protein purification for crystallography and ITC*

After subcloning to pET21a(+), TrpR wildtype and all variants were expressed in *E. coli* BL21(DE3) and purified over Ni-NTA resin and a subsequent Superdex-S75 gel filtration column. All purification steps and measurements were based on the above 50 mM Tris / 300 mM NaCl pH 8 buffer.

#### *Crystallisation, data collection and processing*

Crystals of TrpR wildtype and variants with different ligands were obtained by standard vapour diffusion. The crystals were flash frozen in liquid nitrogen. Data of single crystals were collected at the synchrotron beamline PXII (Swiss Light Source, Villigen PSI, Switzerland) at 100 K and 0.5 degree images were recorded on a Pilatus 6 M detector. Only variant TrpR-M42F-T44L-T81M-N87G-S88Y::IAA was recorded at MX Beamlines BL14.1-3 at BESSY II (Helmholtz-Zentrum Berlin für Materialien und Energie, Berlin, Germany). Data were indexed, integrated and scaled with XDS and converted with XDSCONV[33]. Molecular replacement was performed with Phenix using the coordinates of TrpR-Wt (PDB 1WRP[34] or 1TRO[35]) as search model. Model building was performed with the program Coot[36] and refinement with Phenix[37]. Details on crystallisation conditions, data and refinement statistics for all structures are summarised in Extended data Table S4.

#### *Isothermal Titration Calorimetry (ITC)*

ITC was performed using a VP-ITC (MicroCal). The protein concentration was adjusted to 74µM and 730µM ligand solutions were prepared using the above buffer containing 1% DMSO. Measurements were performed at 20°C with a stirring speed of 300 rpm, reference power 15 µcal/s and spacing of 300 s between injections. The data were analysed using the MicroCal LLC program. Binding data were derived from sigmoidal fits based on a one-site binding model from two measurements for each variant. Heat-of-dilution baselines for the ligands alone were subtracted from the experimental data as described[38].

#### *Test of different FRET pairs*

We tested pairs of Aquamarine, mCerulean3, mTFP1, and mTurquoise2 with Clover, Ypet, and mNeonGreen; Aquamarine additionally with eGFP and mWasabi, mTFP1 with TagRFP. Furthermore, we tested mNeonGreen, Clover, and Ypet with TagRFP and mRuby2. mKO1 was tested with mCherry, mKate2 and mNeonGreen and mWasabi. TagRFP was also tested with mTFP1, mWasabi, mKate2 and mCherry.

#### *Constructs for in vivo AuxSen experiments*

We cloned the final version of the sensor into pJIT6 (pJIT60 2x35S::NLS:AuxSen). Protoplasts were transfected with 10 µg of this construct as previously described[22]. pGIIB pEF-1a::NLS:AuxSen was generated by replacing the Kanamycin cassette of *pGIIK*[39] with *pNos::BAR:tNos* and inserting the sensor between the 1.4 kb promoter of *EF-1a* including the *EF-1a* UTR and *19s* terminator. For expression controls we replaced *AuxSen* by *mNeonGreen* or *GUS*[22]. These constructs were used for transforming plants.
We generated pGIIB pEF-1a::SP:AuxSen:HDEL adding the signal peptide of an Arabidopsis vacuolar basic chitinase and the HDEL ER retention sequence[40] to the above mentioned pGIIB pEF-1a::NLS:AuxSen. This construct was only used in protoplasts.
pGBII UAS::NLS:AuxSen is based on the pGIIB vector, UAS was synthesised by Thermo Fisher Scientific GENEART GmbH, Regensburg, Germany and cloned with the NLS sequence in the *Kpn*I restriction site of the vector and *Bam*HI in front of the sensor. The plants were crossed with RPS5a::Gal4-GR:VP16 UAS:tdTomato. Progeny seedlings were used for the steady-state image acquisitions.

#### *Imaging*

The imaging of the expression controls *pGIIB pEF-1a::NLS:GUS and pGIIB pEF-1a::NLS:mNeongreen* was performed with an Axio Imager Z.1 (Zeiss, Oberkochen, Germany). The GUS staining was done as previously described[22]. Protoplast and seedling images of the AuxSen were acquired with LSM780 (Zeiss, Oberkochen, Germany).

For protoplast imaging of NLS:AuxSen, we excited with 405 nm and detected with 458-498 nm for Aquamarine and 516-621 nm for Neongreen. To detect SP:AuxSen:HDEL we used 405 nm and detected with 458-514 nm for Aquamarine and 517-552 nm for Neongreen, to excite only the Neongreen we used 488 nm. The protoplasts were imaged in K3 solution[41].

For the time series of auxin (and NPA) treatment, root tips of 5 days old seedlings were placed on microscope slides with 10 µM IAA or 50 µM NPA in PBS buffer with 15% glycerol. Regarding the steady-state NPA and BFA images, root segments about 0.1-0.5 cm long were fixed in perfluoroperhydrophenanthrene (PP11; SIGMA-ALDRICH, Saint Louis, USA)[42] and imaged immediately. We excited Aquamarine for FRET with 405, Neongreen with 488 and tdTomato with 561 nm. The emission was collected between 450-490 (Aquamarine), 508-552 (Neongreen) and 600-714 nm (tdTomato).

#### *NPA and BFA treatments*

For the steady-state NPA and BFA experiments, a dexamethasone-inducible two-component system (RPS5A::GAL4-VP16-GR X UAS::NLS:tdTomato) was used. Seedlings were grown as previously described[22], the stratification step varied from 2 to 7 days. The NPA treatment for single acquisition was done placing 5 days old seedlings in ½ MS+S 25 µM DEX plates containing either 50 µM NPA or 0.1 % (v/v) DMSO (control) 24 hours prior to acquisition under the microscope. The procedure for the BFA treatment was the same but with½ MS+S 25 µM DEX plates containing either 10 µM BFA or 0.1 % (v/v) DMSO (control).

#### *Calculation of the FRET ratio*

Image calculations were done with ImageJ Fiji[43,44], the macros used are in the Supplemental Methods. To calculate the FRET ratio we divided the signal intensity of Aquamarine by the signal intensity of mNeongreen. Colours were automatically assigned to range between blue (the nucleus with the minimal FRET ratio) and red (the nucleus with the maximal FRET ratio).

For identification of the nucleus in the roots, the acceptor channel (mNeongreen) was directly excited. For imaging of the dex-inducible auxin sensor, the tdTomato signal was used. For protoplast experiments, the threshold was applied automatically. For measurements with root images, the threshold was done manually with Photoshop. The algorithms in Fiji automatically detected the nuclei from the thresholded images and provided the median FRET ratio for each nucleus. To identify the position of the nucleus with respect to the root, an excel sheet was programmed. Here, the Fiji-extracted nuclei positions were combined with a vector (tip and middle point positions on the root) to obtain the distance of each nucleus to the tip.

### Extended data Figures

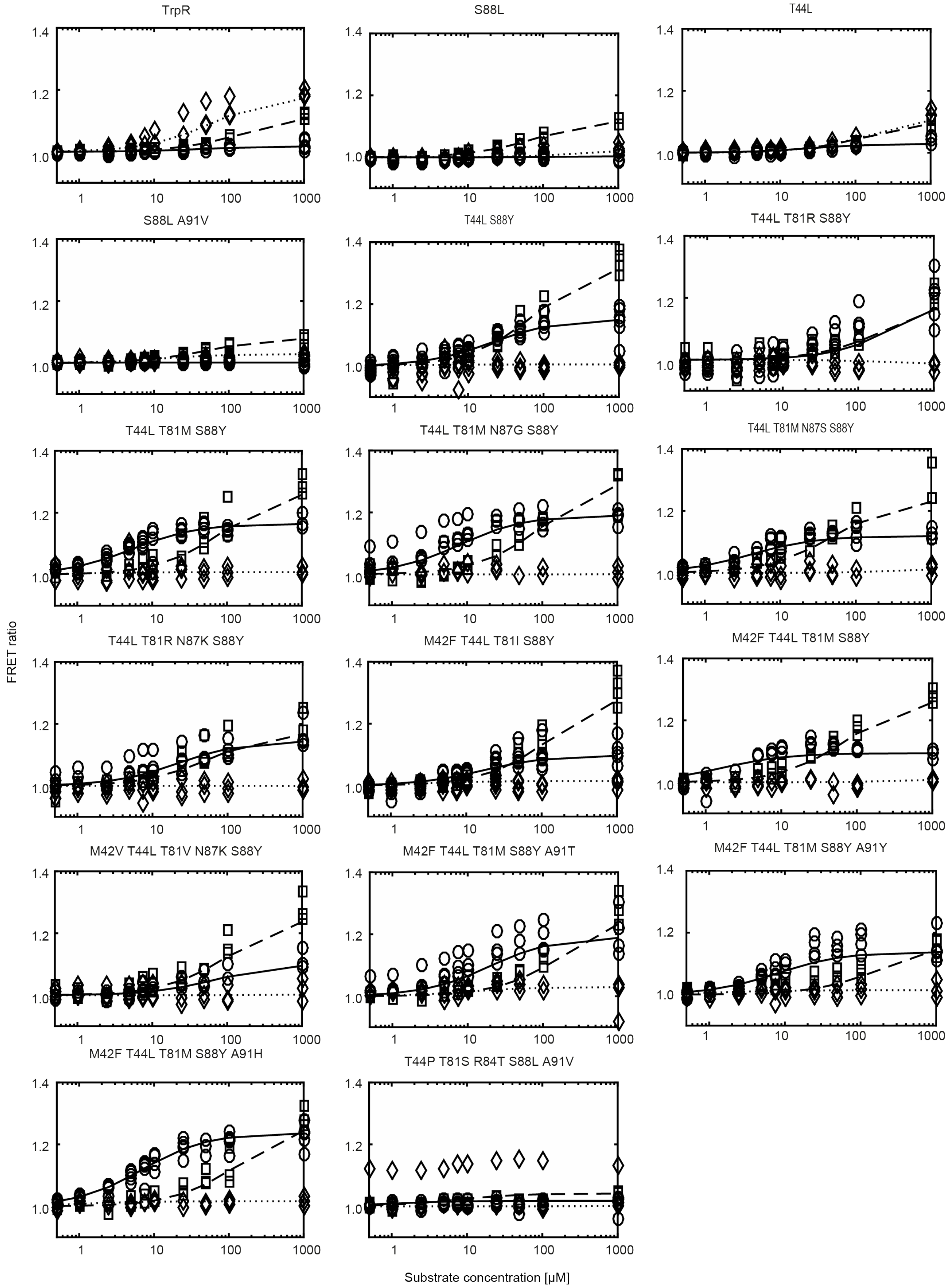

**Extended data Figure S1. FRET response (y-axis) of several binding domain variants to increasing substrate concentration in µM (x-axis).** Circle, IAA; rhombus, TRP; square, IAN.

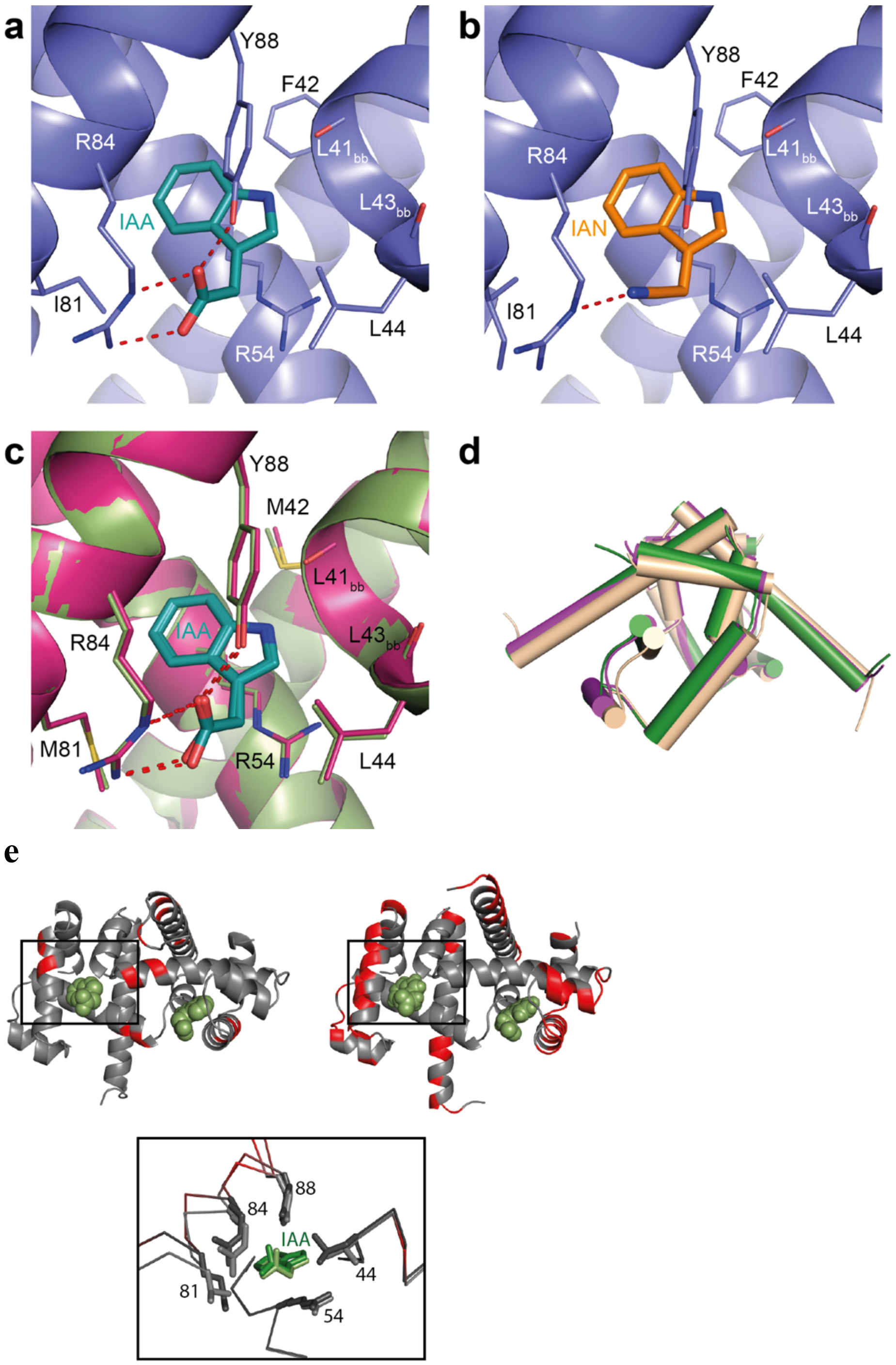

Extended data Figure S2. **Details observed in the crystal structures.** **a**, Structure of IAA in the binding pocket of TrpR-M42F-T44L-T81I-S88Y. **b**, IAN bound to the same variant. **c**, Overlay of IAA in TrpR-T44L-T81M-S88Y (magenta) and TrpR-T44L-T81M-N87G-S88Y (green). **d**, Structural overview of variant TrpR-T44L-T81M-N87G-S88Y (green), TrpR-T44L-T81M-S88Y (magenta), and TrpR-M42F-T44L-T81M-N87G-S88Y (AuxSen, gold). It is apparent that AuxSen differs from the two intermediate structures regarding the overall arrangement of the helices. **e**, Crystal contacts in the tetragonal (P $4_3$, TrpR-S88Y, *left*) and orthorhombic (P $2_1$ $2_1$ $2_1$, TrpR-T44L-T81M-S88Y, *right*) crystal forms. Residues that have symmetry mates within 3 Å distance are shown in red. The ligand IAA is shown in green. The bottom panel shows an overlay of the binding pocket of both structures.

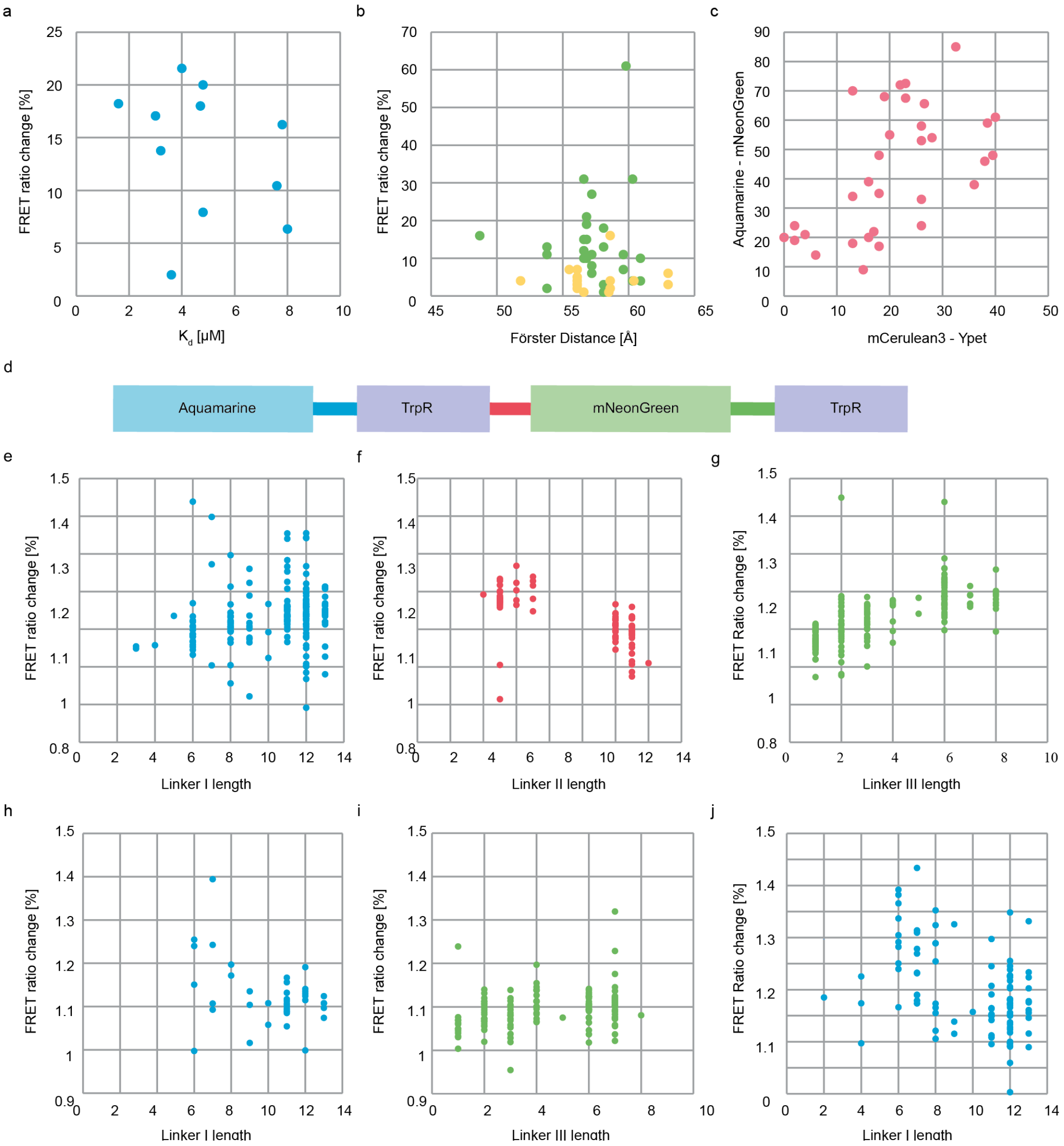

Extended data Figure S3. **Parameters tested for potential influence on FRET ratio change.** **a**, FRET ratio change upon IAA treatment plotted against the Kd of the same variant as determined by ITC. **b**, FRET ratio changes do not correlate with the Förster distance. Blue-yellow pairs are marked in green, yellow-red ones in orange. Note that blue-yellow pairs, in general, show a higher FRET ratio change upon IAA treatment, but a similar range of Förster distances as the yellow-red ones. **c**, FRET ratio changes in [%] of several variants tested with two different fluorophore pairs. Variants showing a strong response with one fluorophore pair usually also show a strong response with another pair (correlation coefficient = 0.6). **d-j**, Effects of mutations in linkers: (**d**) Structure of the construct. The IAA-binding TrpR variants were cloned as tandem repeats into the construct harbouring donor and acceptor fluorophore. (**e-g**) First-round linker mutations. All three linkers were mutated, but no pattern for the optimal linker length could be determined. One linker II variant was chosen for further mutations. (**h-i**) Second-round linker mutations. Linkers I and III were mutated in the variant obtained in the first round, with no changes in the optimised linker II. (**j**) Third-round linker mutations. Linker I was further mutated in the variant harbouring mutations in linkers II and III. The linker length axis indicates the number of amino acid residues.

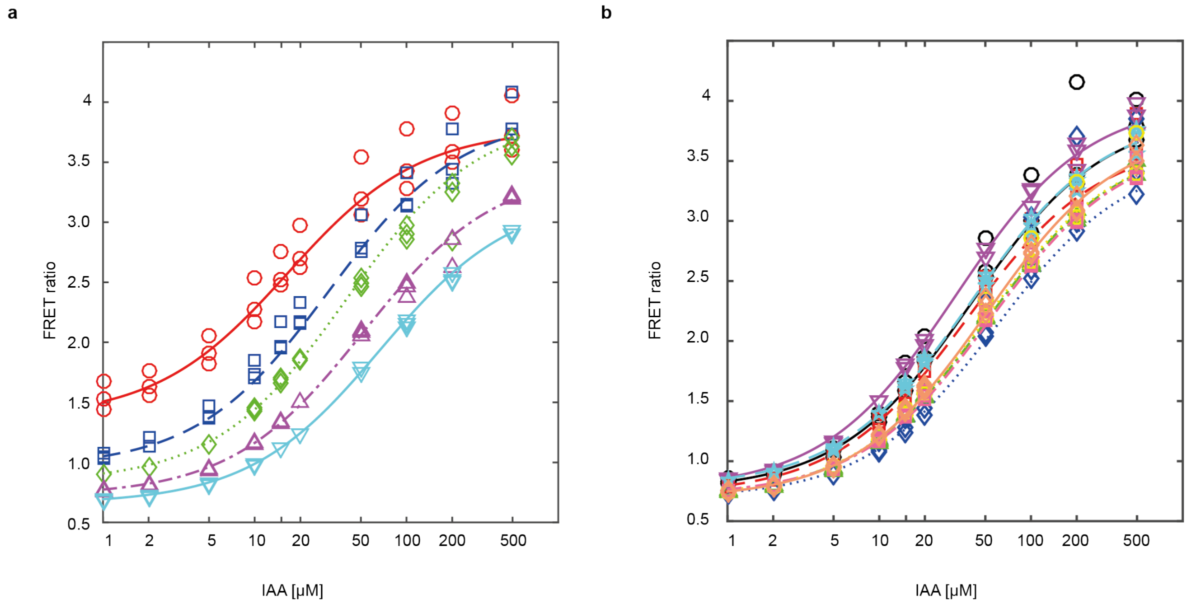

Extended data Figure S4. **pH, salt and redox sensitivity of the auxin sensor.** **a**, The FRET ratio is slightly affected by changes in the pH, but fully functional in the pH range within the plant cell. pH 6.0: red ○, pH 6.5: blue □, pH 7.0: green ◊, pH 7.5: violet Δ, pH 8.0: cyan ∇. **b**, The FRET ratio is not strongly affected by salts and changes in the redox potential. Control, black ○; 1 mM $(NH_4)_2SO_4$, re □; 1 mM $CaCl_2$, blue ◊; 10 mM $NH_4NO_3$, green Δ; 10 mM DTT, violet ∇; 10 mM $H_2O_2$, cyan ☆; 10 mM KCl, yellow ○; 10 mM $KNO_3$, orange □; 10 mM NaCl, orange ◊.

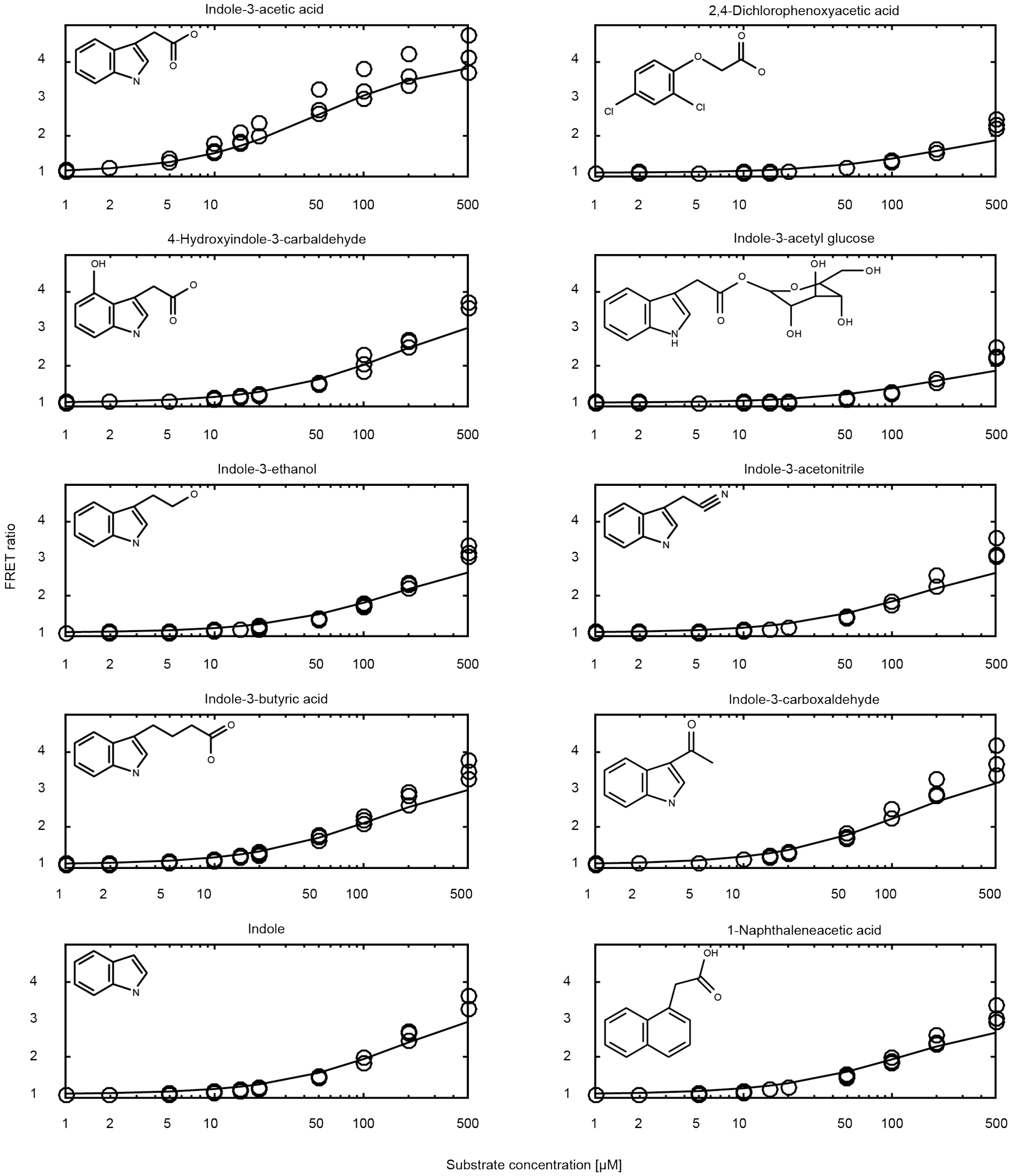

Extended data Figure S5a. **Auxin sensor affinities to auxin-related compounds: some compounds exhibit weak affinities.** The FRET ratio change (y-axis) is plotted against rising concentrations of the individual components in μM (x-axis). Each mark indicates a single measurement.

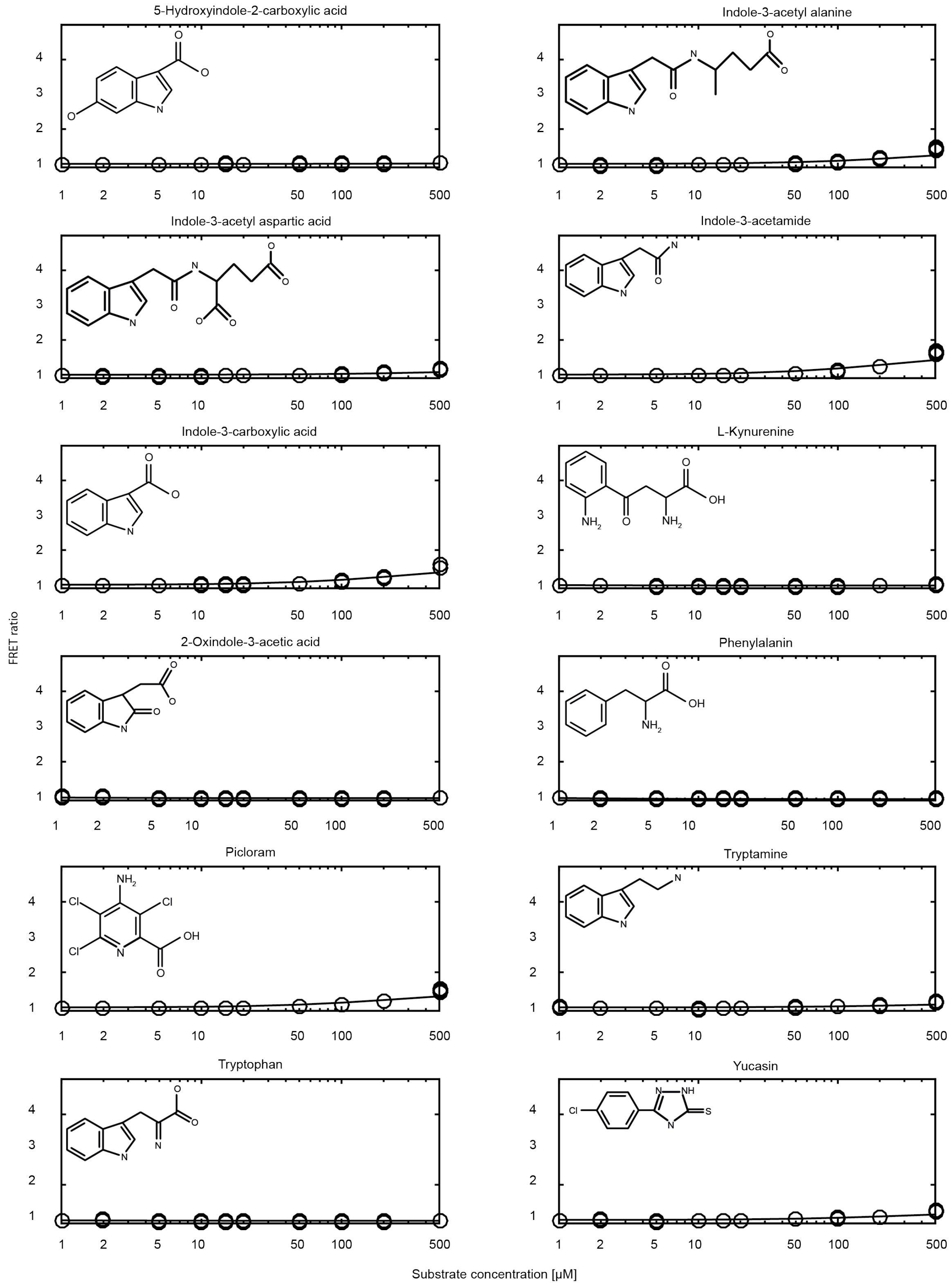

Extended data Figure S5b. **Auxin sensor affinities to auxin-related compounds: several compounds exhibit no affinities.** The FRET ratio change (y-axis) is plotted against raising concentrations of the individual components in µM (x-axis). Each mark indicates a single measurement.

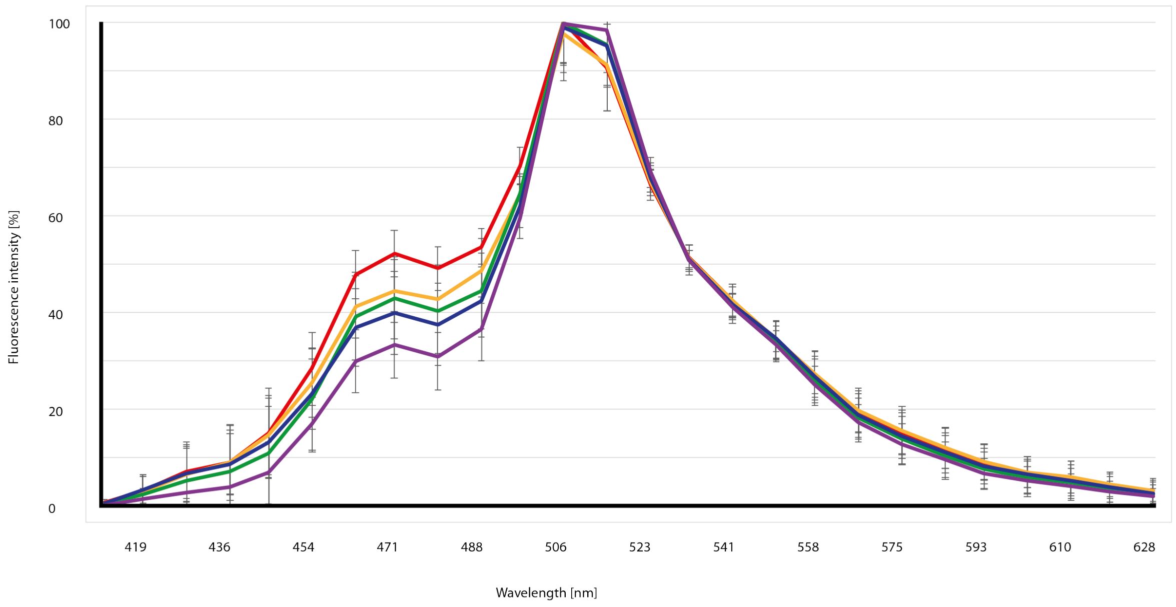

Extended data Figure S6. **Spectra from auxin-treated protoplasts.** Protoplasts transiently expressing the nuclear-localised auxin sensor were incubated in K3 medium with different amounts of IAA: 0 µM, red; 5 µM, orange; 10 µM, green; 20 µM, blue; 50 µM, violet.

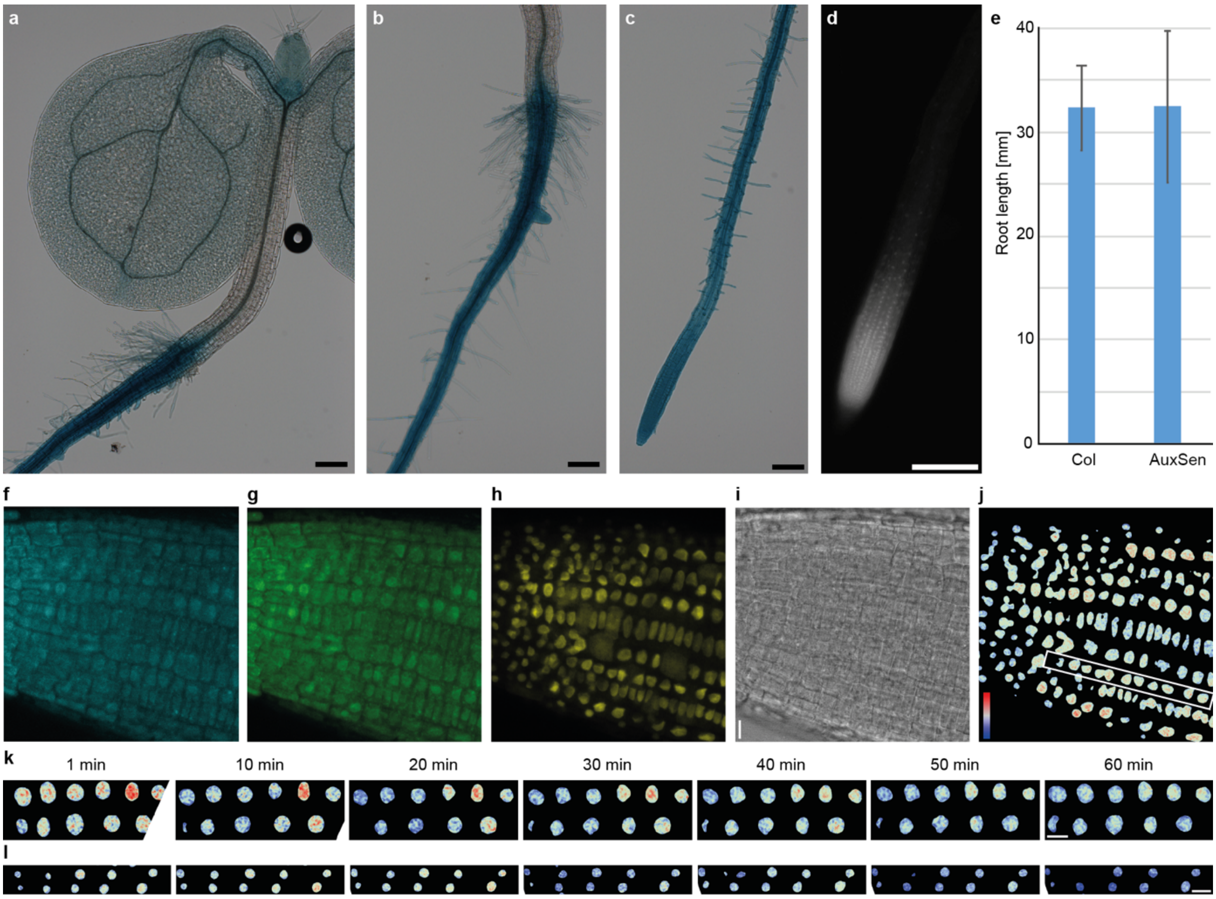

Extended data Figure S7. **EF-1a promoter expression and FRET analysis in the root.** **a-e,** Expression of GUS and mNeongreen under the control of the *EF-1a* promoter. **a-c**, GUS expression. (**a**) The expression is weak in the cotyledons and shoot apical meristem, and absent in the hypocotyl. (**b-c**) Strong GUS expression in the root. **d**, The expression of mNeongreen is especially prominent in the root tip. Scale bar, 200 µm. **e**, No difference in root growth between Col (control) and AuxSen seedlings. n=15; error bars, standard deviation. **f-l,** IAA, IAA and NPA, and control mock treated roots. The root was treated with 10 µM IAA and imaged over 60 minutes. The images show the first time point. **f**, donor signal. **g**, FRET signal. **h**, Direct excitation of acceptor, used for setting the image threshold. **i**, DIC. **j**, Colour-coded FRET ratio with all analysed nuclei; the boxed area is shown in Figure 3c. **k**, The root was pretreated with 50 µM NPA for one hour and then co-treated with 10 µM IAA and 50 µM NPA. **l**, The root was treated with DMSO. Scale bar, 10 µm.

**Extended data Table S1.** In vivo concentration of indole derivatives and induced FRET response of AuxSen.

| Component | FRET ratio fold change at 50 µM | Endogenous concentration | Reference |
|---|---|---|---|
| Indole-3-acetic acid (IAA) | 2.86 | 100 nM | [46, 52, 57] |
| Indole-3-butyric acid (IBA) | 1.7 | <1 nM | [46] |
| 1-Naphthaleneacetic acid (NAA) | 1.48 | - | |
| 2,4-Dichlorophenoxyacetic acid (2,4-D) | 1.13 | - | |
| Indole-3-acetonitrile (IAN) | 1.38 | 100 µM | [46, 21] |
| Indole | 1.45 | 60 nM | [54] |
| Indole-3-carbaldehyde (ICHO) | 1.74 | 100 nM | [50] |
| 4-hydroxyindole-3-carbaldehyde (4-HO-ICHO) | 1.51 | ~300 nM | [50] |
| Indole-3-ethanol (IAEt) | 1.36 | <1 nM | [46] |
| IAA-glucose (Glc) | 1.11 | ~30 nM | [52] |

**Extended data Table S2.** Dissociation constants of selected variants for Trp, IAA, and IAN as determined by ITC.

| Variant | *Ligand* | $K_D$ (μM) | ΔH (kcal mol$^{-1}$) | -TΔS (kcal mol$^{-1}$) | ΔG (kcal mol$^{-1}$) |
|---|---|---|---|---|---|
| **TrpR** | ***Trp*** | 25.0 ± 0.7 | -12.06 ± 0.11 | 5.89 ± 0.12 | -6.18 ± 0.02 |
| | ***IAA*** | 137.4 ± 0.5 | -10.38 ±0.49 | 5.19 ± 0.50 | -5.19 ± 0.01 |
| | ***IAN*** | 63.7 ± 1.7 | -11.76 ± 0.22 | 6.15 ± 0.07 | -5.62 ± 0.01 |
| **TrpR-S88Y** | ***Trp*** | n.d. | n.d. | n.d. | n.d. |
| | ***IAA*** | 14.6 ± 0.1 | -12.62 ± 0.06 | 6.15 ± 0.06 | -6.48 ± 0.01 |
| | ***IAN*** | 17.9 ± 0.6 | -11.62 ± 0.01 | 5.24 ± 0.03 | -6.38 ± 0.03 |
| **TrpR-T44L-S88Y** | ***Trp*** | n.d. | n.d. | n.d. | n.d. |
| | ***IAA*** | 7.8 ± 0.3 | -14.73 ± 0.01 | 7.88 ± 0.03 | -6.85 ± 0.03 |
| | ***IAN*** | 19.5 ± 0.6 | -13.29 ± 0.07 | 6.97 ± 0.09 | -6.32 ± 0.01 |
| **TrpR-T44L-T81M-S88Y** | ***Trp*** | n.a. | n.a. | n.a. | n.a. |
| | ***IAA*** | 3.0 ± 0.1 | -18.75 ± 0.21 | 11.34 ± 0.23 | -7.42 ± 0.02 |
| | ***IAN*** | 33.6 ± 0.2 | -13.55 ± 0.04 | 7.53 ± 0.03 | -6.02 ± 0.00 |
| **TrpR-T44L-T81M-N87G-S88Y** | ***Trp*** | n.a. | n.a. | n.a. | n.a. |
| | ***IAA*** | 1.6 ± 0.0 | -20.60 ± 0.26 | 12.80 ± 0.26 | -7.81 ± 0.02 |
| | ***IAN*** | 46.4 ± 1.7 | -16.31 ± 0.10 | 10.49 ± 0.12 | -5.82 ± 0.03 |
| **TrpR-M42F-T44L-T81M-N87G-S88Y** | ***Trp*** | n.a. | n.a. | n.a. | n.a. |
| | ***IAA*** | 1.5 ± 0.1 | -19.67 ± 0.10 | 11.87 ± 0.12 | -7.82 ± 0.01 |
| | ***IAN*** | 14.3 ± 0.4 | -13.16 ± 0.23 | 6.65 ± 0.21 | -6.51 ± 0.02 |

The dissociation constants were determined via ITC, at 20°C, and pH 8.0. Thermodynamic values for enthalpy (ΔH), entropy (–TΔS), and free energy (ΔG) are shown in kcal mol$^{-1}$.

**Extended data Table S3.** Crystallographic data presented in this work.

**Table S3a** Overview

| Variant | *Ligand* | PDB ID | Resolution | $R_{free}$ / $R_{obs+}$ | Spacegroup | Source |
|---|---|---|---|---|---|---|
| **TrpR, tetragonal** | ***TRP*** | 1ZT9 | 2.0 | 0.219 | P $4_3$ | 1ZT9 |
| **TrpR, orthorhombic** | ***TRP*** | 2OZ9 | 1.7 | 0.180$^{+}$ | P $2_1$ $2_1$ 2 | 2OZ9 |
| **TrpR** | ***IAA*** | 6EJW | 2.0 | 0.243 | P $4_3$ | This work |
| **TrpR-S88Y** | ***IAA*** | 6EJZ | 1.9 | 0.218 | P $4_3$ | This work |
| **TrpR-S88Y-T44L** | ***IAA*** | 6ENI | 1.1 | 0.176 | P $2_1$ $2_1$ $2_1$ | This work |
| **TrpR-T44L-T81M-S88Y** | ***IAA*** | 6EKP | 1.6 | 0.190 | P $2_1$ $2_1$ $2_1$ | This work |
| **TrpR-T44L-T81M-N87G-S88Y** | ***IAA*** | 6ENN | 1.2 | 0.180 | P $2_1$ $2_1$ $2_1$ | This work |
| **TrpR-M42F-T44L-T81M-N87G-S88Y** | ***IAA*** | 6ELB | 1.4 | 0.193 | P $2_1$ $2_1$ $2_1$ | This work |
| **TrpR-M42F-T44L-T81I-S88Y** | ***IAA*** | 6ELF | 1.8 | 0.217 | P $2_1$ $2_1$ $2_1$ | This work |
| **TrpR-M42F-T44L-T81I-S88Y** | ***IAN*** | 6ELG | 1.4 | 0.177 | P $2_1$ $2_1$ $2_1$ | This work |

**Table S3b** Data collection and refinement statistics (molecular replacement) for TrpR-Wt with ligand IAA

| | TrpR-Wt IAA: 6EJW |
|---|---|
| **Data collection** | |
| Space group | $P\ 4_3$ |
| Cell dimensions | |
| $a$, $b$, $c$ (Å) | 81.31 81.31 72 |
| α, β, γ (°) | 90 90 90 |
| Resolution (Å) | 44.93 - 1.991 (2.062 - 1.991) [a] |
| $R_{\mathrm{sym}}$ or $R_{\mathrm{merge}}$ | 0.09365 (1.962) |
| $I / \sigma I$ | 11.55 (0.91) |
| Completeness (%) | 99.45 (96.48) |
| Redundancy | 6.5 (6.0) |
| **Refinement** | |
| Resolution (Å) | 44.93 - 1.991 (2.062 - 1.991) |
| No. reflections | 32088 (3099) |
| $R_{\mathrm{work}}$ / $R_{\mathrm{free}}$ | 0.1945 (0.3088) / 0.2430 (0.3641) |
| No. atoms | |
| Protein | 3293 |
| Ligand/ion | 130 |
| Water | 130 |
| $B$-factors | |
| Protein | 50.17 |
| Ligand/ion | 46.37 |
| Water | 51.45 |
| R.m.s. deviations | |
| Bond lengths (Å) | 0.008 |
| Bond angles (°) | 0.90 |

[a] Numbers in brackets correspond to values in the highest resolution shell.

**Table S3c** Data collection and refinement statistics (molecular replacement) for TrpR-S88Y with ligand IAA

| | TrpR-S88Y IAA: 6EJZ |
|---|---|
| **Data collection** | |
| Space group | P $4_3$ |
| Cell dimensions | |
| *a*, *b*, *c* (Å) | 82.44 82.44 75.65 |
| α, β, γ (°) | 90 90 90 |
| Resolution (Å) | 41.22 - 1.897 (1.965 - 1.897) [a] |
| $R_{sym}$ or $R_{merge}$ | 0.08071 (1.816) |
| $I / \sigma I$ | 20.28 (1.47) |
| Completeness (%) | 99.79 (98.67) |
| Redundancy | 12.8 (12.6) |
| **Refinement** | |
| Resolution (Å) | 41.22 - 1.897 (1.965 - 1.897) |
| No. reflections | 40085 (3931) |
| $R_{work}$ / $R_{free}$ | 0.1840 (0.3045) / 0.2178 (0.3372) |
| No. atoms | |
| Protein | 3337 |
| Ligand/ion | 118 |
| Water | 220 |
| *B*-factors | |
| Protein | 43.80 |
| Ligand/ion | 49.40 |
| Water | 46.76 |
| R.m.s. deviations | |
| Bond lengths (Å) | 0.006 |
| Bond angles (°) | 0.69 |

[a] Numbers in brackets correspond to values in the highest resolution shell.

**Table S3d** Data collection and refinement statistics (molecular replacement) for TrpR-T44L-S88Y with ligand IAA

| | TrpR-T44L-S88Y IAA: 6ENI |
|---|---|
| **Data collection** | |
| Space group | P $2_1$ $2_1$ $2_1$ |
| Cell dimensions | |
| *a*, *b*, *c* (Å) | 54.78 63.41 64.77 |
| α, β, γ (°) | 90 90 90 |
| Resolution (Å) | 34.91 - 1.099 (1.139 - 1.099) [a] |
| $R_{sym}$ or $R_{merge}$ | 0.05169 (0.7009) |
| $I / \sigma I$ | 26.67 (3.85) |
| Completeness (%) | 94.44 (80.40) |
| Redundancy | 12.7 (11.4) |
| **Refinement** | |
| Resolution (Å) | 34.91 - 1.099 (1.139 - 1.099) |
| No. reflections | 87117 (7337) |
| $R_{work}$ / $R_{free}$ | 0.1582 (0.2625) / 0.1755 (0.2627) |
| No. atoms | |
| Protein | 1805 |
| Ligand/ion | 30 |
| Water | 469 |
| *B*-factors | |
| Protein | 14.10 |
| Ligand/ion | 13.10 |
| Water | 23.74 |
| R.m.s. deviations | |
| Bond lengths (Å) | 0.010 |
| Bond angles (°) | 1.08 |

[a] Numbers in brackets correspond to values in the highest resolution shell.

**Table S3e** Data collection and refinement statistics (molecular replacement) for TrpR-T44L-T81M-S88Y with ligand IAA

| | TrpR-T44L-T81M-S88Y IAA: 6EKP |
|---|---|
| **Data collection** | |
| Space group | P $2_1$ $2_1$ $2_1$ |
| Cell dimensions | |
| *a*, *b*, *c* (Å) | 55.035 63.123 64.64 |
| α, β, γ (°) | 90 90 90 |
| Resolution (Å) | 41.48 - 1.458 (1.51 - 1.458) [a] |
| $R_{sym}$ or $R_{merge}$ | 0.03356 (0.7697) |
| $I$ / σ$I$ | 20.30 (1.97) |
| Completeness (%) | 98.54 (94.76) |
| Redundancy | 4.3 (4.2) |
| **Refinement** | |
| Resolution (Å) | 41.48 - 1.458 (1.51 - 1.458) |
| No. reflections | 39383 (3744) |
| $R_{work}$ / $R_{free}$ | 0.1721 (0.3056) / 0.1897 (0.3318) |
| No. atoms | |
| Protein | 1689 |
| Ligand/ion | 41 |
| Water | 250 |
| *B*-factors | |
| Protein | 34.98 |
| Ligand/ion | 30.82 |
| Water | 45.44 |
| R.m.s. deviations | |
| Bond lengths (Å) | 0.002 |
| Bond angles (°) | 0.50 |

[a] Numbers in brackets correspond to values in the highest resolution shell.

**Table S3f** Data collection and refinement statistics (molecular replacement) for TrpR-T44L-T81M-N87G-S88Y with ligand IAA

| | TrpR-T44L-T81M-N87G-S88Y IAA: 6ENN |
|---|---|
| **Data collection** | |
| Space group | P $2_1$ $2_1$ $2_1$ |
| Cell dimensions | |
| *a*, *b*, *c* (Å) | 54.577 63.185 64.904 |
| α, β, γ (°) | 90 90 90 |
| Resolution (Å) | 41.77 - 1.17 (1.212 - 1.17) [a] |
| $R_{sym}$ or $R_{merge}$ | 0.05165 (0.659) |
| $I / \sigma I$ | 16.16 (1.90) |
| Completeness (%) | 98.04 (87.61) |
| Redundancy | 5.8 (4.1) |
| **Refinement** | |
| Resolution (Å) | 41.77 - 1.17 (1.212 - 1.17) |
| No. reflections | 74760 (6616) |
| $R_{work}$ / $R_{free}$ | 0.1483 (0.2250) / 0.1804 (0.2644) |
| No. atoms | |
| Protein | 1837 |
| Ligand/ion | 26 |
| Water | 347 |
| *B*-factors | |
| Protein | 15.68 |
| Ligand/ion | 12.75 |
| Water | 30.90 |
| R.m.s. deviations | |
| Bond lengths (Å) | 0.010 |
| Bond angles (°) | 1.10 |

[a] Numbers in brackets correspond to values in the highest resolution shell.

**Table S3g** Data collection and refinement statistics (molecular replacement) for TrpR-M42F-T44L-T81M-N87G-S88Y with ligand IAA

| | TrpR-M42F-T44L-T81M-N87G-S88Y IAA: 6ELB |
|---|---|
| **Data collection** | |
| Space group | P $2_1$ $2_1$ $2_1$ |
| Cell dimensions | |
| *a*, *b*, *c* (Å) | 55.09 63.31 65.05 |
| α, β, γ (°) | 90 90 90 |
| Resolution (Å) | 45.37 - 1.437 (1.489 - 1.437) [a] |
| $R_{sym}$ or $R_{merge}$ | 0.04813 (1.086) |
| $I$ / σ$I$ | 19.27 (1.37) |
| Completeness (%) | 98.25 (93.32) |
| Redundancy | 7.2 (6.9) |
| **Refinement** | |
| Resolution (Å) | 45.37 - 1.437 (1.489 - 1.437) |
| No. reflections | 41354 (3869) |
| $R_{work}$ / $R_{free}$ | 0.1539 (0.2272) / 0.1931 (0.3159) |
| No. atoms | |
| Protein | 1771 |
| Ligand/ion | 26 |
| Water | 309 |
| *B*-factors | |
| Protein | 24.34 |
| Ligand/ion | 18.70 |
| Water | 36.44 |
| R.m.s. deviations | |
| Bond lengths (Å) | 0.006 |
| Bond angles (°) | 0.82 |

[a] Numbers in brackets correspond to values in the highest resolution shell.

**Table S3h** Data collection and refinement statistics (molecular replacement) for TrpR-M42F-T44L-T81I-S88Y with ligand IAA

| | TrpR- M42F-T44L-T81I-S88Y IAA: 6ELF |
|---|---|
| **Data collection** | |
| Space group | P $2_1$ $2_1$ $2_1$ |
| Cell dimensions | |
| *a*, *b*, *c* (Å) | 54.96 63.41 64.82 |
| α, β, γ (°) | 90 90 90 |
| Resolution (Å) | 31.7 - 1.832 (1.898 - 1.832) [a] |
| $R_{sym}$ or $R_{merge}$ | 0.09423 (1.985) |
| $I / \sigma I$ | 18.08 (1.22) |
| Completeness (%) | 99.50 (96.29) |
| Redundancy | 12.5 (11.2) |
| **Refinement** | |
| Resolution (Å) | 31.7 - 1.832 (1.898 - 1.832) |
| No. reflections | 20416 (1945) |
| $R_{work}$ / $R_{free}$ | 0.1772 (0.4089) / 0.2166 (0.4210) |
| No. atoms | |
| Protein | 1701 |
| Ligand/ion | 26 |
| Water | 155 |
| *B*-factors | |
| Protein | 35.56 |
| Ligand/ion | 31.31 |
| Water | 42.35 |
| R.m.s. deviations | |
| Bond lengths (Å) | 0.013 |
| Bond angles (°) | 1.20 |

[a] Numbers in brackets correspond to values in the highest resolution shell.

**Table S3i** Data collection and refinement statistics (molecular replacement) for TrpR-M42F-T44L-T81I-S88Y with ligand IAN

| | TrpR- M42F-T44L-T81I-S88Y IAN: 6ELG |
|---|---|
| **Data collection** | |
| Space group | $P\ 2_1\ 2_1\ 2_1$ |
| Cell dimensions | |
| $a$, $b$, $c$ (Å) | 55.014 63.468 65.363 |
| α, β, γ (°) | 90 90 90 |
| Resolution (Å) | 42.09 - 1.376 (1.425 - 1.376)[a] |
| $R_{sym}$ or $R_{merge}$ | 0.06254 (0.7719) |
| $I / \sigma I$ | 16.65 (1.83) |
| Completeness (%) | 99.55 (96.20) |
| Redundancy | 6.3 (5.8) |
| **Refinement** | |
| Resolution (Å) | 42.09 - 1.376 (1.425 - 1.376) |
| No. reflections | 47907 (4557) |
| $R_{work}$ / $R_{free}$ | 0.1432 (0.2381) / 0.1765 (0.2949) |
| No. atoms | |
| Protein | 1790 |
| Ligand/ion | 24 |
| Water | 387 |
| $B$-factors | |
| Protein | 16.77 |
| Ligand/ion | 12.77 |
| Water | 29.75 |
| R.m.s. deviations | |
| Bond lengths (Å) | 0.010 |
| Bond angles (°) | 0.99 |

[a] Numbers in brackets correspond to values in the highest resolution shell.
